# The effects of drought and inter-plant competition on the ectomycorrhizal interaction between fungi and Aleppo pine seedlings

**DOI:** 10.1101/2022.10.25.513645

**Authors:** Lior Herol, Hagai Shemesh, Mor Avidar, Shahar Yirmiahu, Yair Zach, Tamir Klein, Stav Livne-Luzon

## Abstract

- An increase in tree mortality is currently evident in forests around the world. Such mortality could be counterbalanced by the native regeneration of seedlings. Seedling establishment under natural conditions is often limited by inter-plant competition and drought conditions. Many forest ecosystems rely on ectomycorrhizal relationships which could be affected by competition and drought, altering forest resilience.
- We carried out an experiment testing the combined effects of drought, herbaceous competition, and ectomycorrhizal fungi (EMF) on the growth and shape of Aleppo pine seedlings and the EMF community composition.
- Pines that germinated in the presence of the EMF spores were taller, had greater biomass, and more side branches. However, under conditions of either competition or drought, the effect of EMF on seedling biomass and height was greatly reduced, while the effect on shoot branching was maintained. Under a combination of drought and competition, EMF had no influence on plant growth and shape. The EMF community was strongly dominated by *Geopora* species, and its structure was not affected by the treatments. Plants experiencing competition were nitrogen poor but presented the highest levels of EMF sequence abundance.
- Stressful conditions seem to alter the relationship between EMF and seedling growth. Specifically, under drought, both colonization and seedling response to EMF was small. However, under competition, colonization was maintained while no growth enhancement was evident. This discrepancy highlights the complexity of the benefits provided to seedlings by EMF under ecologically relevant conditions.

## Introduction

Since the Industrial Revolution, Earth’s climate has been changing at a faster rate than before, resulting in more extreme weather conditions (Höök & Tang, 2013). Some of the obvious outcomes of these changes are increases in dry season temperatures, and longer and more frequent droughts (Brown, 2020). Forests are one of the habitats greatly affected by these climate changes (Change, 2018), expressed in an increase in fire frequency, tree mortality, and slower forest regeneration (Kowaljow *et al*., 2019).

This recent increase in mature tree mortality highlights the role of seedling establishment in guaranteeing the sustainability of forests in the future (Whitmore, 1998). However, seedling establishment is often limited by numerous biotic (Gorchov & Trisel, 2003; De La Cruz *et al*., 2008) and abiotic (Alvarez◻Aquino *et al*., 2004) environmental factors. Because of their small size, seedlings are often at a competitive disadvantage when competing with mature trees (Dickie *et al*., 2005) or herbaceous vegetation (Van Der Waal *et al*., 2009). This disadvantage, which results from their limited access to aboveground (light) and belowground (water and nutrients) resources, can hamper seedling growth and reduce their resilience to droughts (Cavender-Bares & Bazzaz, 2000; Pozner *et al*., 2022) and competition (Fetene, 2003).

The survival, establishment and growth of many tree species depends on mutualistic interactions. Mycorrhiza, the relationship between plants and root inhabiting fungi, is one of the most widespread mutualistic interactions in nature (Peay, 2016). Ectomycorrhiza is a reciprocal relationship in which the tree supplies carbon to the fungi, which in return makes minerals accessible to the tree (Smith & Read, 2010). In addition, the fungi increase the surface area of the root tip, allowing the roots to absorb minerals and water from a wider surface (Wu *et al*., 2012). Moreover, for many tree species, the ectomycorrhizal relationship is obligatory (Smith & Read, 2010). Therefore, ectomycorrhiza is a mutual interaction that greatly affects the health of the forest and its persistence (Rudgers *et al*., 2007).

The nature of mutualistic interactions can depend on environmental conditions (Begon *et al*., 1986). In the case of ectomycorrhiza, benign conditions, such as high nutrient availability (Bai *et al*., 2020), have been shown to limit fungal colonization of roots. This is probably the result of independent resource uptake by plant roots, making the interaction redundant from the plant’s perspective. However, natural conditions are often not benign, and mutualism can also be limited by competition over shared resources or by the ability of one or both partners to cope with environmental stress. It is therefore expected that under natural conditions, mutualisms in general and ectomycorrhiza specifically, would range from being mutually beneficial to being detrimental (Johnson *et al*., 1997).

The interaction between drought and EM symbiosis, did receive some attention in the literature (Garbaye, 2000; Lehto & Zwiazek, 2011; Nickel *et al*., 2018; Sebastiana *et al*., 2018; Wang *et al*., 2021). Specifically, under drought conditions, mycorrhizae were found to transfer water between trees (Egerton-Warburton *et al*., 2007; Kakouridis *et al*., 2020). Moreover, ectomycorrhizal mycelium can benefit trees by extracting inaccessible water from micro-crevasses within soil particles (Bornyasz *et al*., 2005). Wang et al. (2021) showed that under water limited conditions, ectomycorrhizal fungi can alleviate hydraulic failure and reduce carbon starvation of plants, which leads to an increase in plant growth while reducing their mortality rate. However, all these plant benefits are dependent on the EMF ability to survive and perform under stress. Because both the fungi and the plant require water, it is not clear whether under drought conditions the ectomycorrhizal interaction would remain beneficial for the plant, or rather shift to neutrality, or even become detrimental to the plant.

While drought is a pivotal stressor for seedlings in many ecological systems, it is often combined and possibly enhanced by competition with herbaceous vegetation which associate with arbuscular mycorrhizal fungi (AMF). While the role of AMF in inter-plant competition has received considerable attention (Scheublin *et al*., 2007; Facelli *et al*., 2010; Klironomos *et al*., 2011; Lin *et al*., 2015; Tedersoo *et al*., 2020) the influence of EMF on a seedling’s ability to cope with other plant competitors was hardly studied. The most direct way EMF can assist tree seedlings experiencing competition is by improving its uptake of soil resources, specifically nutrients (Marschner & Dell, 1994; Chalot *et al*., 2002; Smith & Read, 2008). However, although the biology of EMF nutrient uptake has been extensively studied, it was practically unstudied in the setting of inter-plant competition. We are aware of only 4 studies that tested of the effect of EMF on inter-plant competition. Specifically, EMF was found to alleviate the negative effects of inter-specific competition between seedlings of EM trees (Perry *et al*., 1989; Shi *et al*., 2017), as well as increase phosphorus uptake of pine seedlings competing with a grass (Pedersen *et al*., 1999) and growth of pine seedlings competing with an AM perennial (Peay, 2018).

To the best of our knowledge, no manipulative studies have been conducted on the interaction between drought, inter-plant competition and EMF in their effect on seedling establishment. In this study we tested whether EMF provide an advantage to Allepo pine seedlings experiencing drought, competition with an annual grass, or both. We hypothesized that the relative advantage that the EMF can provide to the pine seedlings will increase under drought and/or competitive conditions since both stressors will minimize the ability of the plant to acquire soil resources independently.

## Methods

### Experiment overview

Mediterranean forests are prone to relatively frequent environmental changes (Petit *et al*., 2005). The effects of these changes, which are often the result of fires (Glassman *et al*., 2016) or droughts (García de Jalón *et al*., 2020), might be mitigated by the ectomycorrhizal fungal community that had evolved in these habitats. Aleppo pine (*Pinus halepensis* Miller) is the most common forest tree species around the Mediterranean (Ne’eman & Osem, 2021) and is well adapted to local conditions (Klein *et al*., 2011; Voltas *et al*., 2018; Patsiou *et al*., 2020). This species is also well known for its mycorrhizal connections, including many EMF species (Avital *et al*., 2022; Cahanovitc *et al*., 2022).

To test our hypothesis, we tested the effect of the ectomycorrhizal fungi (EMF) on the performance (biomass and shoot branching) of Aleppo pine seedlings under ectomycorrhizal presence (with/without), water conditions (full irrigation/ drought) and competition (with/without). The complete crossing of these three factors created 8 different groups. Each group was replicated 28 times (2 water ×2 EMF×2 competition × 28 replications = 224 pots). Six spare plants were added as an incomplete block resulting in a total of 230 pots.

### Growth conditions

The experiment was conducted under natural daylight (12-13 hours) and annual ambient air temperatures (21.66±6 °C) in a net-house in Tel-Hai College, Israel (N 35°34’41"E 11°33’14”). The experiment was initiated during February 23-27, 2020 and was conducted for six months.

### Potting material

Grassland soil and sand were used as potting material. The soil was collected in herbaceous Mediterranean grassland, lacking EM host plants - according to previous experiments (Livne◻Luzon *et al*., 2017; Livne-Luzon *et al*., 2021b) from a field near Tel-Hai College (N 35°34’47"E 58°33’14”). The potting material was mixed (50% sand, 50% grassland soil) using an electric cement mixer. The pots (4 liters) were filled with the mixture.

### EMF

Forest soil was collected from four different locations (Table S1). All forest soils were collected underneath Aleppo pine trees. At every location, soil was sampled at four locations spaced 1-2 meters apart. The depth of the taken soil was a few centimeters below the surface in order not to collect the remnants of the organic matter. All the soils collected were mixed and sieved (2mm). The forest soil (100 ml) was mixed into the EMF treatment pots. To maintain an equal volume, 100 ml of potting material (50% sand, 50% grassland soil) was mixed into the pots without the EMF. In order to increase the study’s ecological validity, we chose to add natural forest soil which includes a diverse EMF community (Livne◻Luzon *et al*., 2017; Avital *et al*., 2022). This decision limits the internal validity of the study by increasing the number of alternative explanations regarding the obtained results. We find the effect of other soil microbiota in the inoculum to be the main concern. However, we interpret the change in root allocation (a known effect of EMF; Smith & Read, 2010) and the effect of inoculation on EMF read abundance as strong indicators of EMF as the main drivers of the discovered effects. In light of the above, we find natural inoculum to be preferred over a species poor synthetic community.

### Plant material

Pinecones were collected three days before the beginning of the experiment, at the first location (“Road view” near Kfar Gilady, Table S1). The pinecones were placed in an oven to release the seeds from the cones (15 min, at 80 C°). Four seeds were sown in each pot, and the pots were arranged in blocks (8 pots per block). All the pots were irrigated with a computerized irrigation system (20 minutes in the morning every day, at a capacity of one liter per hour). On the 8^th^ of April 2020, seedlings were thinned down to one per pot.

### Competition stress

On February 27^th^ 2020, *Hordeum Spontaneous* seeds were sown in the pots experiencing competition (4 in every pot). On 8^th^ April 2020, *Hordeum Spontaneous* were thinned down to one per pot. *Hordeum Spontaneous* (K. Koch) is a fast-growing grass common in the southern and eastern Mediterranean.

### Drought conditions

On the 30^th^ of April 2020, the pots in the drought treatment were removed from the irrigation system. The pots were irrigated manually to create drought stress (Table S2). The weight of the water in field capacity was calculated by reducing the weight of the dry pot (the pot’s mass after drying in the oven - 4.375 kg), from the weight of a pot at field capacity (5.75 kg). The difference is the weight of the water in field capacity (1.375 kg). In the drought treatment, the pots were maintained at 10% volumetric capacity (0.1375+4.375=4.52). The amount and frequency of the irrigation was decided upon in accordance with the average weight of the pots. The changes in the weight of the pot were caused because of changes in the weather during the experiment as well as plant growth. Ten pots were weighed prior to every irrigation event and 40 pots (10 pots from each treatment) were weighed four times during the experiment (Table S2).

### Morphological measurements

On August 6, 2020 the plants’ height and branches were measured. The purpose of these measurements was to characterize morphological differences between the different treatments. We standardized the branch number by dividing the number of branches by the total plant biomass. This was done to obtain an index of the number of branches, irrespective of the size of the plant.

### Harvesting protocol

On August 23-25 2020, plants were harvested by block. Plants were removed intact from the pots and washed gently under tap water and the roots and shoots were separated. Roots were scanned visually for colonized root-tips and all colonized root tips were removed using sterilized forceps, inserted into a 1.5 ml Eppendorf tube added with 300 μl CTAB buffer, and stored in a −20°C freezer until DNA extraction. To avoid cross-contamination between samples, all tools were sterilized using ethanol (70%).

Both roots and shoots were placed separately in the oven (60 C°, for 3 days). Shoot and root mass were measured using an analytical scale (Radwag, AS 220.R2, Radom, Poland). Out of the initial 230 plants, 17 plants died during the experiment and 2 plants had extreme biomass values (4.45g, z=3.75; 2.52g, z=4.41) and have been excluded (with no qualitative changes to the results; Fig S1, Table S3) from further morphological analyses.

### Molecular identification of fungal species

DNA was extracted from 76 random root samples (see number of samples in Table S4) following the methods of Livne-Luzon et al., (2017). Briefly, frozen root tips were bead beaten (at least 2 × 30 seconds at 4000 rounds per minute till fine powder was achieved), and DNA was extracted from each root tip sample following a modified version of the QIAGEN (Valencia, CA, USA) DNA easy Blood and Tissue Kit. Barcoded amplicon sequencing of the fungal ITS2 region was performed on a MiSeq platform (Illumina, San Diego, CA, USA). A two step protocol for library preparation was performed according to Straussman lab (Nejman *et al*., 2020), with several modifications; First PCR reactions were performed using KAPA HiFi HotStart ReadyMix DNA polymerase (Hoffmann-La Roch, Basel, Switzerland) in 50 μl reaction volumes with 5 μl DNA extract, and 1 μl of every primer 5.8S-Fun (5′-AACTTTYRRCAAYGGATCWCT) (Taylor *et al*., 2016) and RD2-ITS4Fun (5′AGACGTGTGCTCTTCCGATCT-AGCCTCCGCTTATTGATATGCTTAART). The reverse primer consisted of the ITS4-Fun primer (Taylor *et al*., 2016) with the linker adapter RD2. PCR reactions were performed as follows: initial 2 min at 98 °C followed by 35 cycles of 10 s 98 °C, 15 s 55 °C, and 35 s 72 °C final cycle with 5 min 72 °C. Second PCR reactions were performed using the same DNA polymerase in 50 μl reaction volumes with 1/10 (5 μl) of the first PCR reaction, and 1 μl of every primer P5-rd1-5.8S-Fun (5′-AATGATACGGCGACCACCGAGATCT-ACACTCTTTCCCTACACGACGCTCTTCCGATCT-AACTTTYRRCAAYGGATCWCT) and RD2-Barcode (5′ AGACGTGTGCTCTTCCGATCT-BARCODE). The forward primer consisted of the adaptor p5 the linker RD1 and the primer 5.8S-Fun. The reverse primer consisted of the adapter RD2 and the individual barcode. PCR reactions were performed similar to the first PCR but with only 6 cycles. PCRs were cleaned using Qiaquick PCR purification kit (Qiagen, Hilden, Germany), quantified fluorescently with the Qubit dsDNA HS kit (Life Technologies Inc., Gaithersburg, MD, USA). Libraries were quality checked for concentration and amplicon size using the Agilent 2100 Bioanalyzer (Agilent Technologies, Santa Clara, CA, USA) and size selected with AMPure magnetic beads (Beckman Coulter Inc., Brea, CA, USA). We sequenced all the samples in one amplicon using Illumina MiSeq technology with 300 bp paired-end reads (PE300_V3) in the Grand Israel National Center for Personalized Medicine (Weizmann institute of science, Rehovot, Israel).

### Bioinformatics

We used R (R Core Team, 2018, version 4.0.3) and the R-Studio for bioinformatics and statistical analysis. Raw sequences were demultiplexed and adapters together with barcodes were removed for 76 root samples (Water × Ectomycorrhiza × Competition). The sequences were analyzed using the amplicon sequencing dada2 package v. 1.7.9 in R (Callahan *et al*., 2016). In summary, sequences were quality-filtered and trimmed. We only used sequences longer than 50 bases with a mean number of expected errors below 2 (maxN = 0, maxEE = c(2,5) minLen = 50 truncQ = 2). Paired-end sequences were merged using the MergePairs function. We then applied a dereplication procedure on each sample independently, using derepFastq function. Finally, all files were combined in one single Fasta file to obtain a single amplicon sequence variant (ASV) data file. We removed singletons (minuniquesize = 2) and de novo chimera sequences using removeBimeraDenovo function against the reference database (UNITE/UCHIME reference datasets v.7.2). Sequences were then clustered, and taxonomic assignment (id = 0.98) was done against the UNITE database. Non-fungal ASVs were removed. FUNguild was then used to parse ASVs into ecological guilds (Nguyen *et al*., 2016); we then filtered the table to include only highly probable EMF genera. Due to the low colonization rates of the young seedlings, the average sequence abundance was very low (the mean sequence abundance per sample was 1142±1516, for *Geopora* 1167±1507) and the total number of putative EMF ASVs (16) was low as well.

### Shoot nitrogen quantification

During the analysis of the biomass and genetic results we developed a post-hoc hypothesis regarding the role of competition for nitrogen (see discussion). We then decided to quantify the nitrogen content of the pine shoots. Dry pine needles were used to quantify the shoot’s nitrogen content. The nitrogen content was estimated only for five experimental blocks (a total of 40 plants). In order to choose representative blocks, standard Z scores were calculated for the mean total plant biomass of every block. The 5 blocks with the smallest absolute Z scores were used in the analysis. Two hundred mg of needles were taken from each seedling. In small plants with less than 200 mg, all needles were used (156.6 ± 49mg for all plants combined). Nitrogen content was quantified following the Kjeldahl method using Kjeltec™◻ 8100 (Foss, Denemark). Nitrogen content was then divided by the needle biomass to attain the percent of nitrogen in the tissue.

### Statistical analysis

We used general linear mixed models, using a fully factorial design, to account for differences in the plant’s total, shoot and root dry biomasses, plant height, branch density and the relative abundance of the most dominant EMF taxa (i.e., *Geopora)*. The following explanatory variables were included as fixed factors: EMF inoculation, water and competition treatments as well as their interactions. The experimental blocks were included in the model while allowing for random slopes and intercepts for each fixed factor. Due to high variation and skewedness of the data, the branch density, and the sequence abundance of *Geopora* were log transformed prior to analysis. Figures were generated using R packages ggplot2 (version 3.3). Illustrations were created with BioRender.com

## Results

For a comprehensive details regarding the statistics see table 1.

**Table 1:**
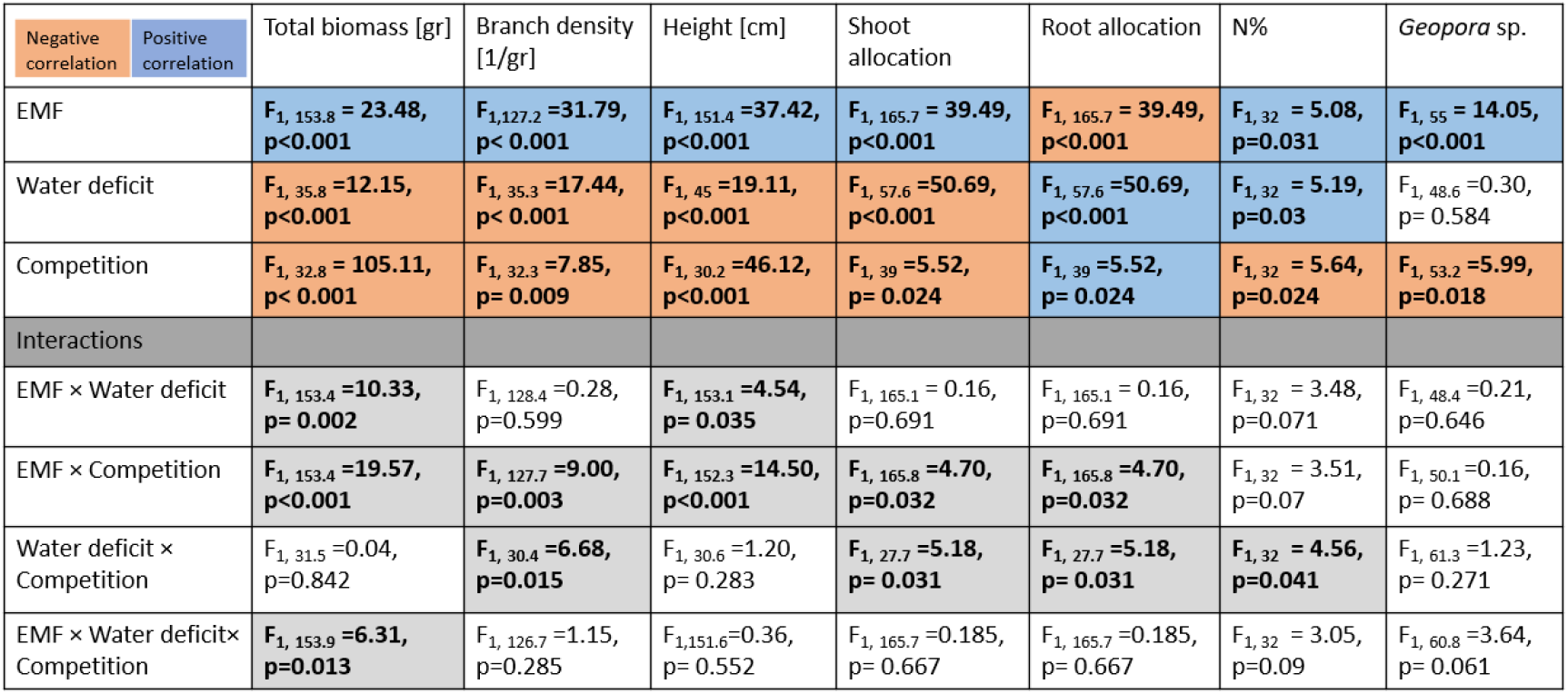
statistical analysis of plant growth and genetic analysis according to water, competition and ectomycorrhiza treatments and the interactions between them. The orange color represents a negative connection between the treatments and the growth index/ EMF present, while the blue color represents a positive connection.

### Plant response to treatments

As expected, drought caused a decrease of 20% to the plant’s total biomass (Control: 0.588±0.442; Water: 0.47±0.288; F_1,35.8_ =12.15, p<0.001, Fig.1a). Similarly, competition caused a decrease of 60% in the plant’s total biomass (Control: 0.736±0.399; Competition: 0.297±0.16; F_1, 32.8_ = 105.11, p< 0.001, Fig.1a). However, ectomycorrhizal inoculation led to an increase of 34% in plant biomass (Control: 0.456±0.218; EMF: 0.61±0.479; F_1, 153.8_ = 23.48, p<0.001, Fig.1a). Nevertheless, the positive effect of ectomycorrhiza was more evident when the plants were watered frequently and the soil was wetter (Water × Ectomycorrhiza: F_1, 153.4_ =10.33, p=0.002, Fig.1a). In addition, the positive effect of ectomycorrhiza was more evident when the plants were not facing competition (Competition × Ectomycorrhiza: F_1, 153.4_ =10.33, p<0.001, Fig.1a). Interestingly, when plants were not experiencing any stress, ectomycorrhiza had a positive effect on the total biomass of the plants. However, when the plants were experiencing either one stress or the combined stress of both drought and competition, ectomycorrhiza did not increase the plants’ total biomass (Water × Ectomycorrhiza × Competition: F_1, 153.9_ =6.31, p=0.013, Fig. 1a).

**Fig.1.**
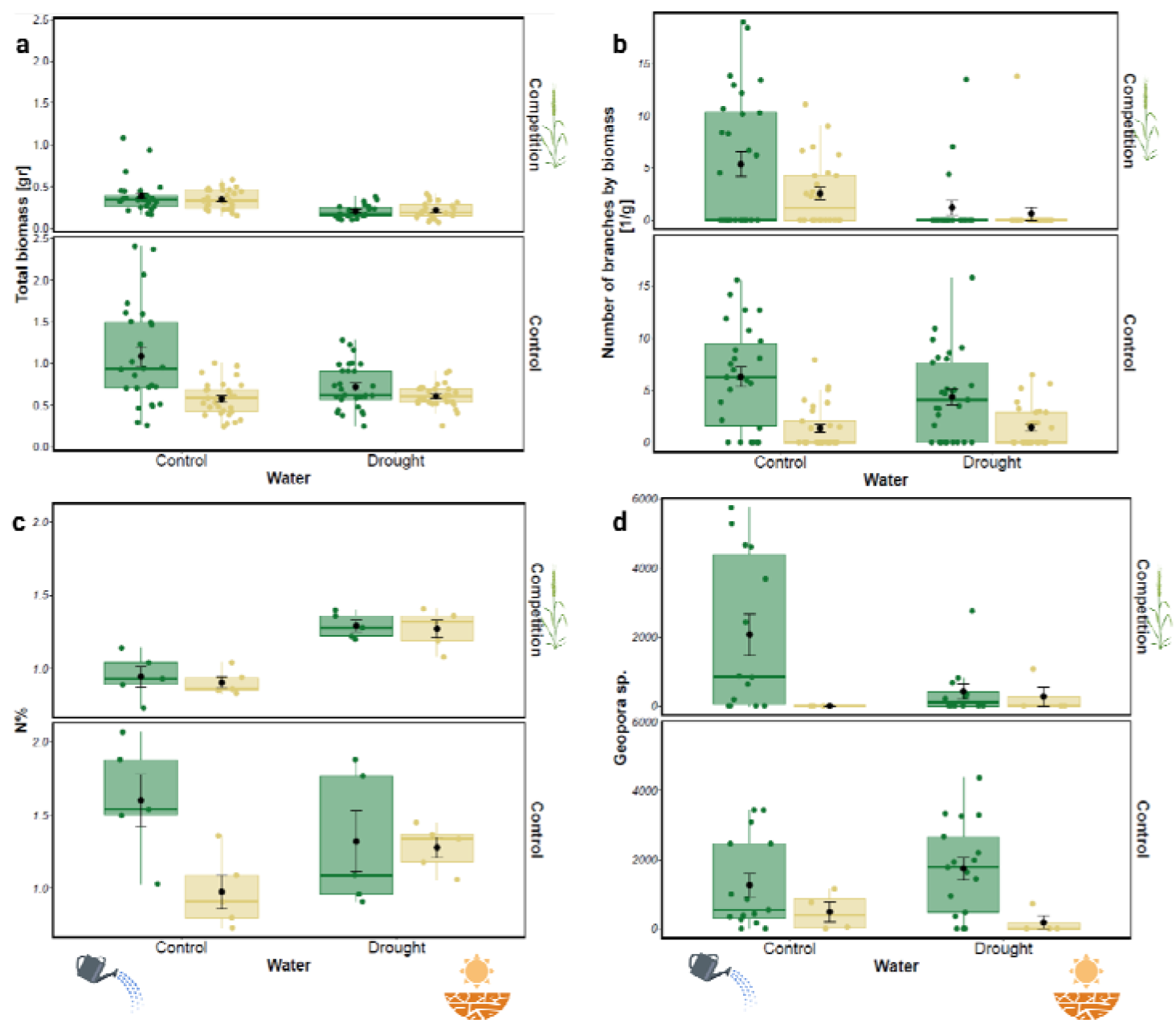
Pine seedling (a) total biomass [g], (b) number of branches by biomass [1/g], (c) needles N% and (d) *Geopora* sp. sequence abundance, according to water, competition and ectomycorrhiza treatments. The first and third hinge of each box plot represent the 25th and 75th percentile, the Like the plant’s total biomass, ectomycorrhizal inoculation led to an increase of 192% to the plants’ branch density (Control: 1.52±2.64; EMF: 4.44±5.14; F_1,127.2_ =31.79, p< 0.001, Fig.1b), while drought and competition both caused a decrease of 46% and 19% to the plants branch density (Control: 3.83±4.84; Water: 2.06±3.49; F_1, 35.3_ =17.44, p<0.001, and Control: 3.27±4.0; Competition: 2.66±4.7; F_1, 32.3_ =7.85, p=0.009, for drought and competition respectively; Fig.1b). However, competition had a stronger negative effect than drought (Water × Competition: F_1,30.4_ =6.68, p=0.015, Fig.1b). In addition, the positive effect of ectomycorrhiza was more evident when the plants were not experiencing competition (Ectomycorrhiza × Competition: F_1,127.7_ =9.0, p=0.003, Fig.1b).

As expected, drought and competition both caused a decrease of 15% and 25% to the plant’s height (Control: 8.17±3.07; Water: 6.91±2.36; F_1, 45_ =19.11, p<0.001, and Control: 8.57±3.03; Competition: 6.43±2.04; F_1, 30.2_=46.12, p<0.001 for water and competition respectively, Fig. S2). Furthermore, ectomycorrhizal inoculation led to an increase of 27% in plant height (Control: 6.66±1.6; EMF: 8.48±3.43; F_1, 151_ =37.42, p<0.001, Fig.S2). However, the positive effect of ectomycorrhiza was more evident when the plants were either watered frequently (Water × Ectomycorrhiza: F_1, 153.1_ =4.54, p= 0.035, Fig. S2) or not experiencing competition (Ectomycorrhiza × Competition: F_1, 152.3_ =14.5, p< 0.001, Fig.S2).

Similarly, drought caused a decrease of 12% to the plant’s shoot allocation (Control: 0.66±0.09; Water: 0.58±0.08; F_1,57.6_ =50.69, p<0.001, Fig. S3), while competition and ectomycorrhizal inoculation led to an increase of 5% and 11% to the plant’s shoot allocation (Control: 0.60±0.09; Competition: 0.34±0.09; F_1,39_ =5.52, p=0.024, and Control: 0.59±0.08; EMF: 0.65±0.09; F_1, 165.7_ = 39.49, p<0.001 for competition and EMF respectively, Fig. S3). However, the negative effect of competition was more evident when the plants were under water stress (Water × Competition: F_1, 27.7_ =5.18, p= 0.031, Fig.S3). In addition, the positive effect of ectomycorrhiza was more evident when the plants were not experiencing competition (Ectomycorrhiza × Competition: F_1,165.8_ =4.7, p=0.032, Fig. S3). Similar yet opposite results were obtained for the plant’s root allocation.

### Pine nitrogen content

As expected, competition caused a decrease of 15% to the plant’s nitrogen content (Control: 1.3±0.39; Competition: 1.1±0.21; F_1,32_ =5.64, p=0.024, Fig.1c). Drought caused an increase of 14% in the plant’s nitrogen content (Control: 1.11±0.375; Water: 1.29±0.24; F_1,32_ = 5.19, p=0.03, Fig.1c). Ectomycorrhizal inoculation also led to an increase of 14% in plant nitrogen content (Control: 1.11±0.23; EMF: 1.29±0.38; F_1,32_ = 5.08, p=0.031, Fig.1c). Nevertheless, the negative effect of competition was only evident when water was not limiting (Water × Competition: F_1, 32_ =4.56, p=0.041, Fig.1c).

### Ectomycorrhizal fungal colonization

As expected, during root tip collection we did not observe colonized root-tips on plants which were not amended with EMF inoculum. However, some levels of colonization were revealed during the molecular sequencing. Overall, in the inoculated plants, molecular identification of EMF root-tips yielded a relatively short list of EMF species. These species mostly belong to two main taxa, namely, *Geopora* and *Tuber*. The most dominant genus was *Geopora* (99.44%), which had a higher sequence abundance on the roots of plants inoculated with EMF in comparison to non-inoculated plants (F_1, 55_ =14.05, p<0.001, Fig.1d). On average, the sequence abundance of *Geopora* decreased in the presence of a competitor (F_1, 53.2_ =5.99, p=0.018, Fig.1d). However, this was evident only under drought (**non-significant**; Ectomycorrhiza ×Water × Competition: F_1, 60.8_ =3.64, p= 0.061, Fig.1d), it appears that when plants were suffering a single stress, *Geopora* was still present on their roots. However, when encountering the combined stress of both drought and competition, *Geopora* sequence abundance was reduced to the same level as in plants that were not inoculated at all. In addition, there was a significant positive correlation between branch density and sequence abundance of *Geopora* (Pearson’s r = 0.438, p<0.001).

## Discussion

Droughts, which are expected to become more frequent due to climate change, are considered to be one of the largest threats to forested habitats. In addition to lack of water, drought conditions can increase inter-plant competition (Kaisermann *et al*., 2017) which influences the establishment of tree seedlings. We hypothesized that ectomycorrhizal fungi would provide an advantage to pine seedlings in general and more so under stressful conditions. Our hypotheses were not fully supported by the data. Specifically, while forest soil inoculum resulted in increased growth under all conditions, its positive effect was greatly diminished (becoming not significant) under either, each or both stresses. Surprisingly, the shoot branching pattern of the seedlings showed a qualitatively different response to inoculation compared to those of the seedling height and mass.

Although the interaction between plant and EMF is considered mutualistic, we found a discrepancy between EMF presence and their effect on seedling growth. While EMF were present and beneficial under benign conditions (Rincón *et al*., 2007), under each of the single stresses, EMF colonization (estimated from sequence abundance) was maintained, although no apparent advantage was observed regarding plant growth (Fig. 1a, d). The preservation of the interaction even when it does not seem to benefit plant growth is somewhat surprising. This apparently altruistic pattern can be the result of unmeasured benefits such as pathogen resistance (Gonthier *et al*., 2019) or the plant lack of ability to reject the fungal symbiont. The fact that the fungi sustain on the roots under harsh conditions could also provide an advantage in case conditions improve in the future. Furthermore, inconsistent with our hypothesis, under the combined stress, the seedlings demonstrated both reduced growth and a reduction in EMF sequence abundance. We interpret this to mean that under single stress conditions, the fungi can survive but are unable to provide growth benefits to the seedlings. When both stressors occur simultaneously, it seems as if the fungi’s mere survival is hampered.

Our results indicate that when water was not limited, competition stress still had a negative effect on seedling growth (Fig. 1a) and nitrogen content (Fig. 1c). However, under these conditions, fungal abundance was highest (Fig. 1d). It is well established that low levels of nutrients (especially nitrogen) encourage EMF colonization (Smith & Read, 2010). The high EMF abundance found in the competition treatment when water was not limiting, could be the result of inter-plant competition over nutrients. It therefore seems that the plant needs the fungi the most when minerals are limited, however, the fungi can colonize the roots only when its own basic requirements for soil moisture are met (Fig. 2).

**Fig. 2.**
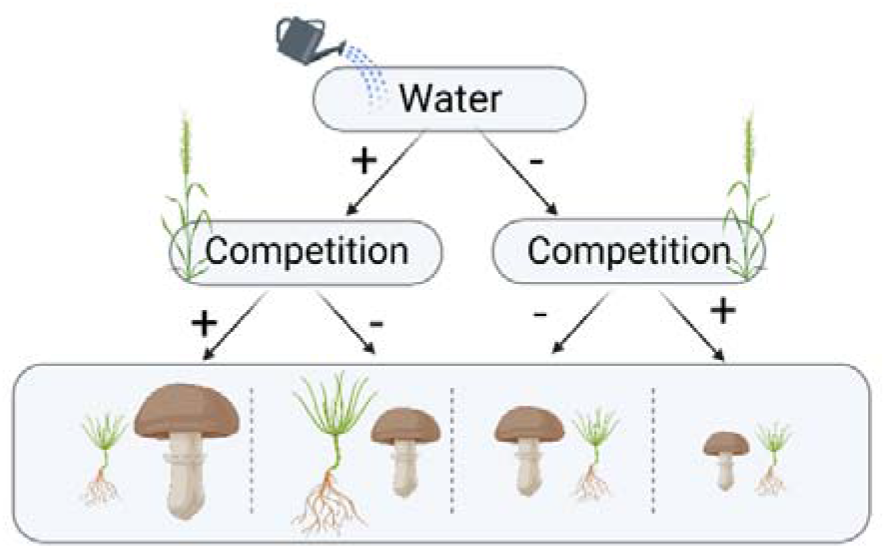
A conceptual hierarchy of the effect of both water and competition stress on the EM relationship. The size of the fungi and the seedling represent the approximate amount of Geopora sp. sequence abundance and the seedlings total biomass in the water/competition treatments.

The fact that under competition and ample watering EMF did not have a positive effect on plant biomass (Fig. 1a), suggests that water is a prerequsite for fungal survival but it not sufficient to enhance plant growth. This complexity might be related to the soil organic matter available for EM decomposition. We hypothesize that the available inorganic minerals in the soil were quickly uptaken by the grass while the EMF were limited in their ability to asist the pine seedling due to the limited size of the pot and the organic material content of the potting material used in this study. Future studies manipulating EMF, competition and soil organic matter under greenhouse and field conditions could shed more light on this complex interaction.

Unlike the effects of ectomycorrhiza on biomass, which disappeared under a single stress, the effect on shoot branching was maintained under either competition or drought but not under both (Fig. 1b). This positive effect on shoot branching was evident even after controlling for plant biomass. In certain ecological scenarios, more shoot branching can be advantageous to plants under competitive conditions (Evers *et al*., 2011), however, this can greatly depend on the competing plant community, light availability and the species biology and strategies (Gruntman *et al*., 2017). The mechanism linking EMF colonization with shoot branching is currently unknown. A similar effect was found in redwood seedlings inoculated with AMF (Willing, 2019) and in a previous study with Aleppo pine seedlings (Livne-Luzon *et al*., 2021a).

Interestingly, while the growth effects of EMF diminished under a single stress, the shoot branching effect was maintained and was correlated with fungal abundance. Therefore, the mechanism linking EMF and shoot branching seems to function independently of the known growth effects of EMF. We suggest that the shoot branching mechanism is dependent on the mere presence of the fungi, which might induce a hormonal effect on the shoot branching pattern. For example, production of the growth hormone auxin was connected with EMF colonization in Poplar (Felten *et al*., 2009), and cypress saplings exuded auxin when inoculated with rhizosphere bacteria (Oppenheimer-Shaanan *et al*., 2022). Moreover, an increase in ABA hormone, which is related to drought tolerance, was seen in redwood seedlings colonized by AMF under drought (Willing, 2019). This proposed mechanism should be tested under different ecological scenarios, to better understand the role of the fungi in changing the shoot structure and the consequences of this change to plant fitness.

In conclusion, when environmental conditions are benign, EMF contributes to plant growth. However, under single or combined stressors, EMF presence did not contribute to the plants’ total biomass. Nonetheless, when plants were experiencing a single stress, EMF presence still increased their branch number. Although plant growth was similarly reduced by both single stresses, drought seems to reduce the presence of EMF while competition seems to increase it. We assume that minimal water availability is necessary for both plant functioning and fungal survival. Furthermore, plants experiencing drought might be limited in their ability to support their mycorrhizal partners. On the other hand, plants experiencing competition rely on their mycorrhizal partners to decompose organic matter and alleviate nutrient stress caused by their competitors and should be more engaged with EMF. The role of EMF in elevating drought and competition stresses of establishing seedlings seems to be complex and further studies, especially about inter-plant competition, are needed. Deciphering the role of EMF in seedling establishment under drought and competition could help us better predict forest dynamics under current global changes.

## Acknowledgements

We wish to thank Rog Ido and Shifra Avital, for their help with setting-up the illumine library, to Dener Efrat for her assistance with the statistical analysis and to Yvonne Lipman for the English editing.

