## Supplementary information for "The effects of drought and inter-plant competition on the ectomycorrhizal interaction between fungi and Aleppo pine seedlings"

Table S1: Forest soil locations

|  | Location |
| --- | --- |
| (N 35°33'49"E 59°33'13") | Road view" near Kfar Giladi |
| (N 35°33'34"E 24°33'14") | "The roaring lion", Kfar Giladi |
| (N 35°34'24"E 46°33'14") | The surrounding road of Kfar Giladi |
| "(N 35°33'07"E 29°33'13") | Forest near "Mitzpe Adi |

Table S2: Manual irrigation details

| Average of the weight of the pots | Amount of water (ml) | Date |
| --- | --- | --- |
| 4.72±0.24 | 400 | 7.5.2020 |
| 4.75±0.44 | 400 | 12.5.2020 |
| 4.637±0.21 | 400 | 14.5.2020 |
| 4.61±0.22 | 400 | 18.5.2020 |
| 4.92±1.02 | 400 | 21.5.2020 |
| 4.66±0.2 | 400 | 26.5.2020 |
| 4.7±0.21 | 400 | 31.5.2020 |
| 4.69±0.22 | 400 | 4.6.2020 |
| 4.66±0.2 | 400 | 8.6.2020 |
| 4.73±0.22 | 400 | 14.6.2020 |
| 4.69±0.21 | 200 | 18.6.2020 |
| 4.65±0.2 | 400 | 23.6.2020 |
| 4.7±0.23 | 200 | 28.6.2020 |
| 4.69±0.23 | 200 | 1.7.2020 |
| 4.67±0.23 | 200 | 5.7.2020 |
| 4.7±0.24 | 200 | 9.7.2020 |
| 4.71±0.23 | 200 | 13.7.2020 |
| 4.72±0.24 | 200 | 17.7.2020 |
| 4.68±0.23 | 200 | 22.7.2020 |
| 4.67±0.23 | 200 | 26.7.2020 |
| 4.69±0.22 | 100 | 2.8.2020 |
| 4.65±0.22 | 100 | 6.8.2020 |
| 4.62±0.21 | 200 | 11.8.2020 |
| Average= 4.69±0.06 | Average= 286.96±111.53 | Average time gap= 4.36±1.02 |

Table S3: Statistical analysis of plant growth and genetic analysis according to water, competition and ectomycorrhiza treatments and the interactions between them with outliers. The orange color represents a negative connection between the treatments and the growth index/ EMF present, while the blue color represents a positive connection.

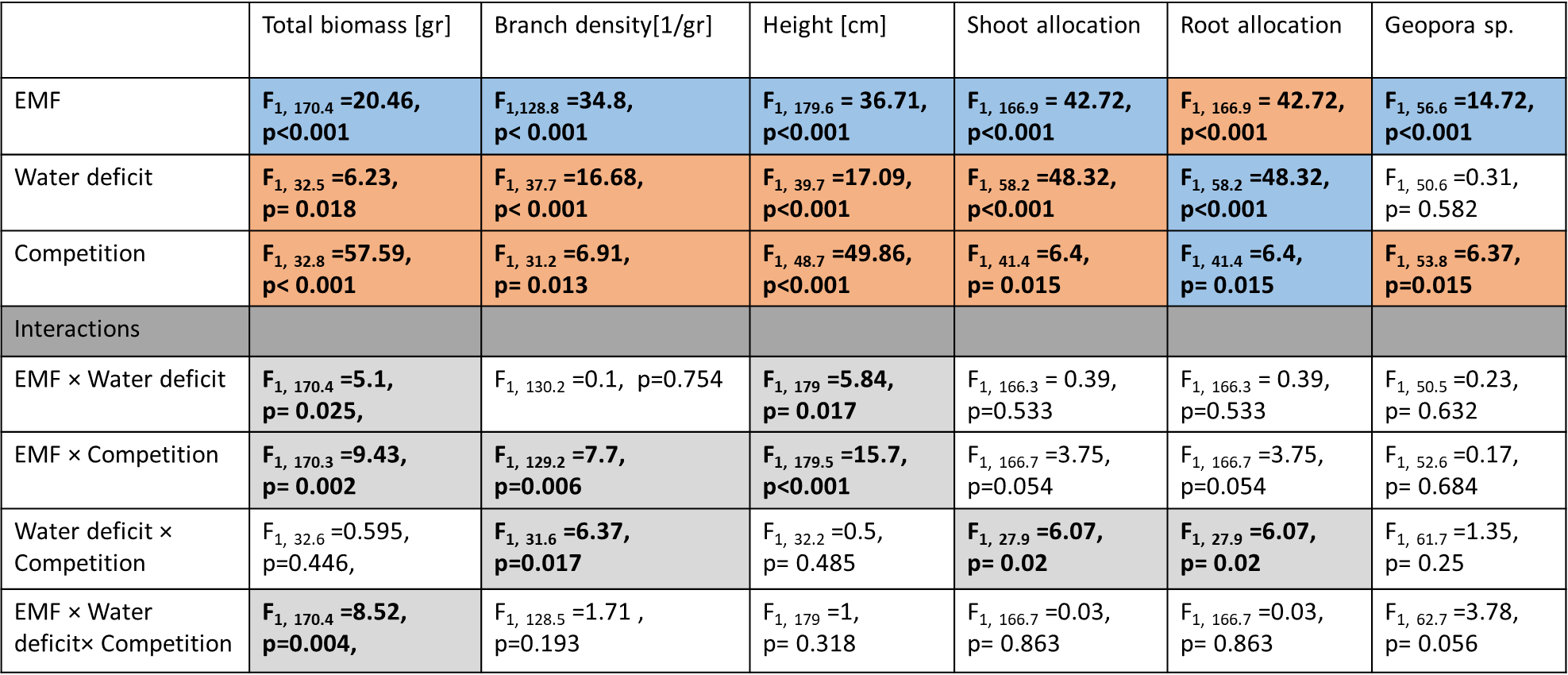

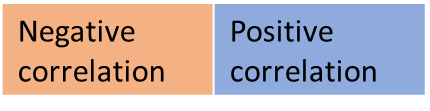

Table S4: Number of samples for molecular identification of fungal species‏

| - | + | - | + | Competition  \  Drought |
| --- | --- | --- | --- | --- |
| 17  Drought stress | 13  Double stress | 4  Drought stress | 4  Double stress | + |
| 16  Optimal condition | 14  Competition stress | 4  Optimal condition | 4  Competition stress | - |
| With EMF presence | With EMF presence | No EMF presence | No EMF presence | EMF |

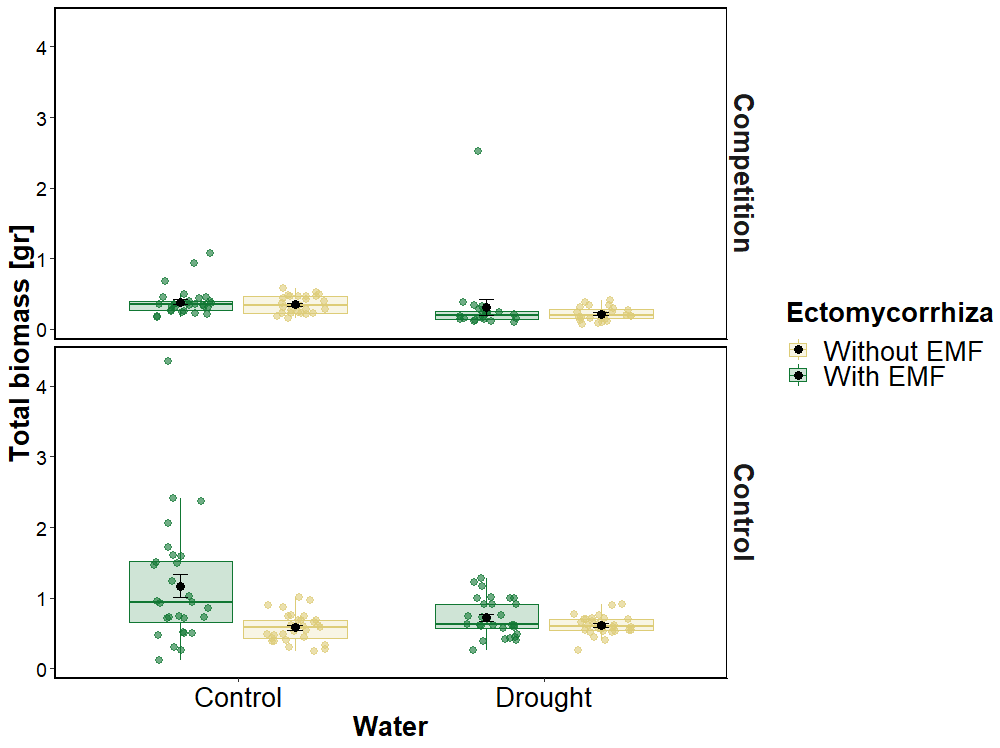

Fig.S1 Pine seedling total biomass [g] according to water, competition and ectomycorrhiza treatments with outliers. The first and third hinge of each box plot represent the 25th and 75th percentile, the middle hinge is the median and the black point is the mean ±SE

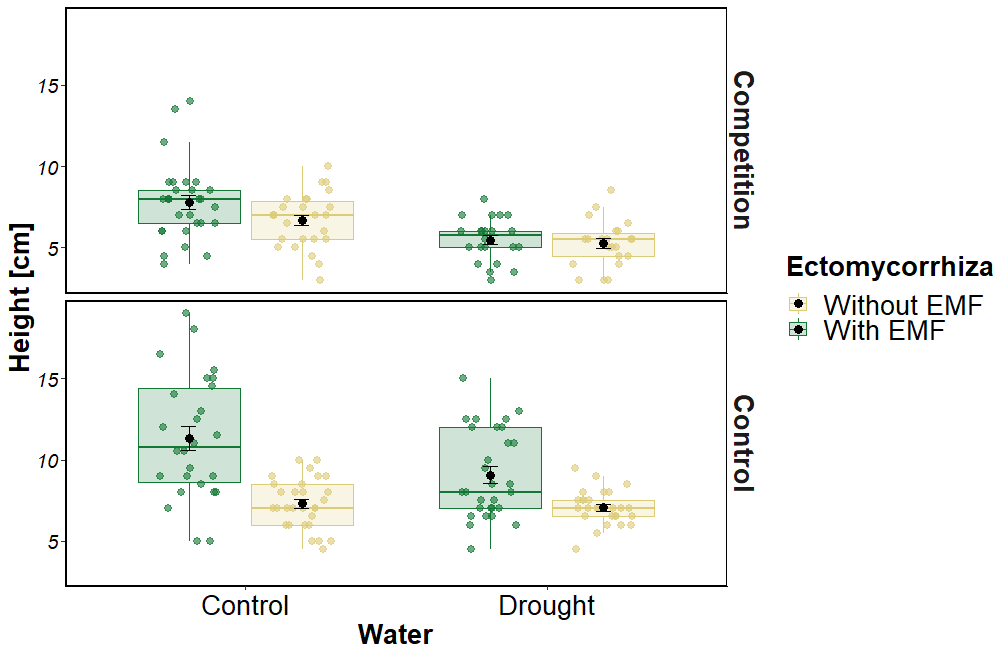

Fig.S2 Pine seedling height according to water, competition and ectomycorrhiza treatments. The first and third hinge of each box plot represent the 25th and 75th percentile, the middle hinge is the median and the black point is the mean ±SE

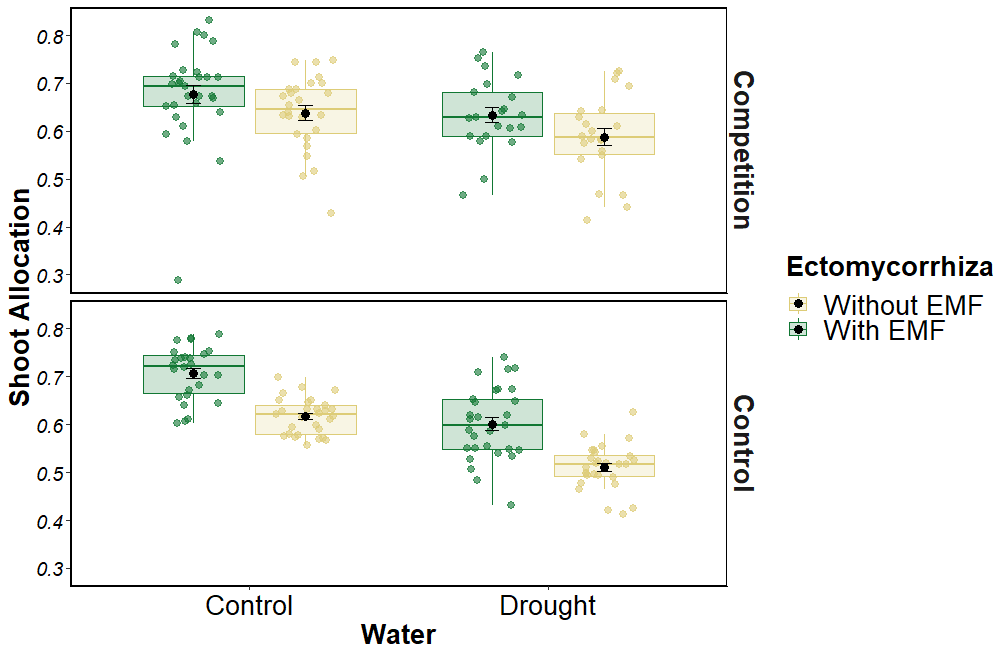

Fig.S3 Pine seedling shoot allocation according to water, competition and ectomycorrhiza treatments. The first and third hinge of each box plot represent the 25th and 75th percentile, the middle hinge is the median and the black point is the mean ±SE

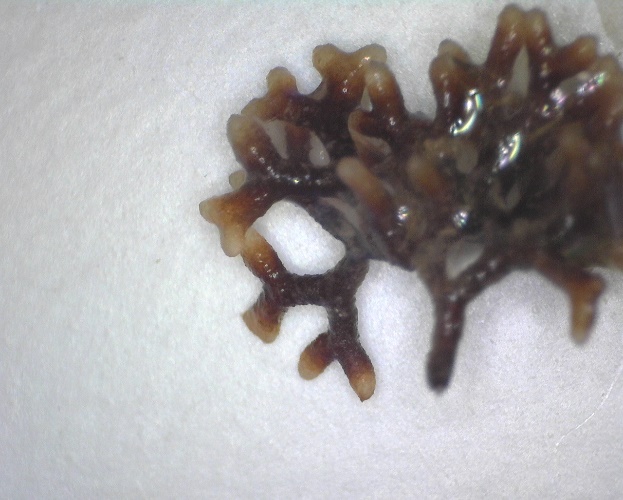

Fig. S4: typical mycorrhizal root tip morphology later identified as colonized by *Geopora* sp.
